# The All Window-Size Search method for improved statistical power in multiple comparisons correction

**DOI:** 10.64898/2026.07.14.738000

**Authors:** Matthew Nelson

## Abstract

Correcting for multiple comparisons is a fundamental challenge throughout the biological sciences, particularly for data sampled over ordered continua such as time, space, or frequency. Existing approaches, including cluster-based permutation tests and threshold-free cluster enhancement (TFCE), leverage spatial or temporal contiguity but remain dependent on predefined statistical frameworks or thresholding procedures. Here we introduce the All Window-Size Search (AWSS) method, a permutation-based procedure that formally controls the family-wise error rate while adaptively searching across all contiguous window sizes and locations. For each permutation, test statistics are summed across every possible window, generating null distributions of maximal statistics at every window size. A second stage estimates the null distribution of the most significant uncorrected p-value that would arise from searching across all window sizes, allowing final p-values to be corrected for the adaptive search process itself. This procedure statistically formalizes the implicit multiscale search that investigators naturally perform when visually inspecting ordered data. Simulations with known ground-truth effects demonstrate that AWSS can provide substantially greater statistical power than conventional cluster-based permutation methods for broad, low-amplitude effects while maintaining appropriate family-wise error control. Because the framework is independent of any particular statistical test, it is readily applicable to diverse forms of one-dimensional ordered data. Here we test this application with simulations as well as using real human sEEG neural recording data. Future extensions will generalize the method to multidimensional spatial and spatiotemporal datasets, including neuroimaging and other high-dimensional biological data.

## Introduction

The multiple comparisons statistical problem affects nearly every researcher. It poses particular problems for datasets involving data that is discretely sampled over a continuum when there is not knowledge of the precise group of samples within that domain that are expected to be statistically significant. In neuroimaging, the problem presents itself, for example, when a researcher analyzes fMRI voxels across an entire brain (consisting of tens of thousands of voxels) without knowing which voxels potentially show an effect. If one imagines a case where the data was pure noise, 5% of the voxels will be found to be “significant”, based on the simple definition of the null hypothesis of the statistical test. Thus the hypothesis test performed on each voxel needs to be appropriately correct for the family-wise error rate of tests performed, however the standard procedure of Bonferroni correction is non-viable because of the large number of hypothesis tests to correct across. Every study must appropriately address this problem; consideration of it is not optional.

Multiple strategies have been developed to address this problem. False discovery rate (FDR)(Benjamini & Hochberg, 1995) procedures control the expected proportion of false positives across tests and are widely used in neuroimaging and electrophysiology; however, they treat individual samples independently and do not leverage contiguity of effects across samples. Cluster-based permutation approaches(Maris & Oostenveld, 2007) partially address spatial or temporal dependence by identifying clusters of supra-threshold samples and evaluating cluster-level statistics under permutation. While effective in many settings, cluster-based methods depend on the choice of a cluster-forming threshold and do not explicitly correct for adaptive selection of cluster extent. Threshold-free cluster enhancement (TFCE) was developed to reduce reliance on arbitrary thresholding by integrating across cluster-forming thresholds(Smith & Nichols, 2009); however, it still operates within a predefined enhancement framework and does not directly model the distribution of the best effect across all contiguous window sizes. Thus, despite important advances, a formal statistical framework that explicitly corrects the adaptive multi-scale search process remains lacking.

## Methods

The method. The All Window-Size Search (AWSS) method is illustrated graphically in **Figure 1**. It is shown here for the simple application of investigating the existence and locations of regions of significant differences between two curves. First, in order to generate the necessary null distributions, the experimental condition assigned to each recorded data sample (e.g. a trial) are randomly shuffled nShuffs times.

**Figure 1.**
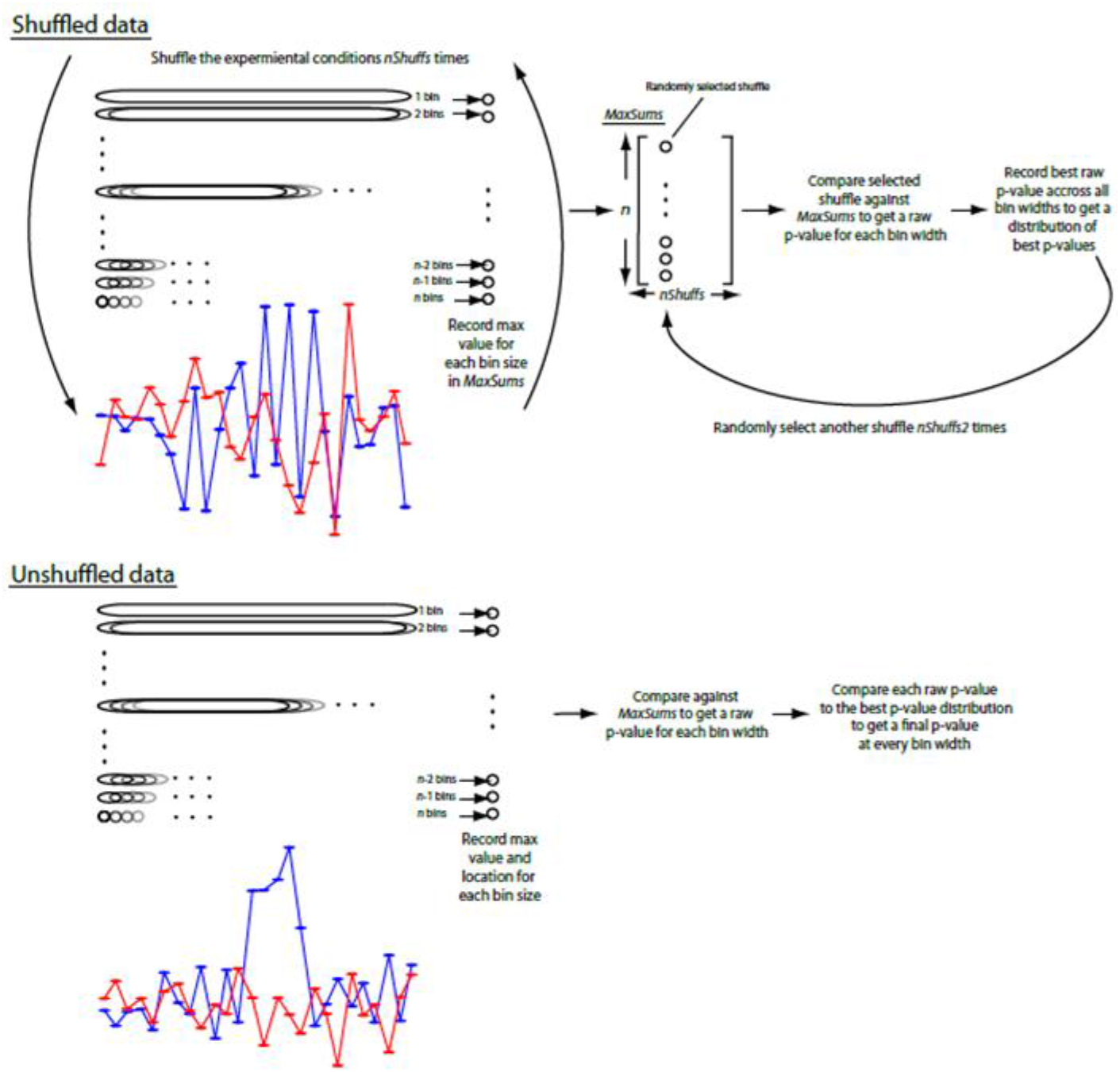
Schematic Illustrating the all bin-width search method to uncover the locations of significant differences between two data curves. The two hypothetical data curves shown in the upper left and lower left panels would reflect the mean curves over several measurements of the curve (e.g. trials). Upper left panel: The experimental conditions between the two curves are randomly assigned to each trial. The test statistic (e.g. the mean difference between the curves) is summed across adjacent samples for each possible location at each possible bin width. The maximum sum for every bin width for each shuffle is recorded in the array MaxSums. Upper right panel: Individual shuffles from MaxSums are randomly selected and compared against the rest of MaxSums to get a raw p-value at each band width. The best p-value across band-widths for each selected shuffle is recorded to provide a distribution of best raw p-values. Lower left panel: The location and value of the maximum sums for every bin width in the unshuffled data is recorded one time. Lower right panel: The maximum sums are compared against MaxSums to get a raw p-value at every bin width. These raw p-values are then compared against the best p-value distribution to get a final p-value at every bin width.

For each shuffle, the test statistic is calculated at every sample point in the data. In this case, the test statistic is simply the mean difference between the two (shuffled) conditions. Adjacent test statistics are summed across adjacent time points/locations over all possible window sizes and locations. For data of length n samples, at a window size of 1 there will be a test statistic at all n possible locations, at a window size of 2 there will be a summed test statistic over two adjacent samples at all n-1 possible locations, proceeding all the way to a bin size of n for which there will be only one test statistic at the only possible location, i.e. summed over all the data samples. For each shuffle, the maximum value for every bin-width is recorded in the n x nShuffs sized array MaxSums.

After completing this process nShuffs times, the array MaxSums can then be used as a null distribution across all nShuffs shuffles of the raw, unprotected p-value for the summed test statistics at every window size. If we had pre-specified a particular bin-width, we could stop here, and use the corresponding distribution of test-statistics in MaxSums as a null-distribution to compare the results of the unshuffled data. Note that it would it would not be appropriate to look at the data, then choose one particular window-size to test, and test it in such a manner. One’s eyes will automatically perform such a test across all window sizes anyway.

However, to perform an analysis at an unspecified bin-width, one needs to investigate the distribution of unprotected p-values across bin-widths, and provide an estimate of how often in randomized data that by chance the best p-value across all bin-widths will reach a certain level. To achieve this, an individual shuffle within MaxSums is randomly selected nShuffs2 times. Each randomly selected shuffle is compared against the distribution of shuffles in MaxSums to provide a raw, unprotected p-value at every bin-width. This is given by the proportion of shuffles in MaxSums with a value of the best sum at a given bin-width greater than or equal to the value for the chosen shuffle at the corresponding bin width. This raw p-value represents the probability that the given sum would be as high for the given bin-width by chance. Then, the best (minimum) raw p-value across all bin-widths for that shuffle is recorded. This process is repeated nShuffs2 times, resulting in a distribution of the best raw pvalues present in randomized data. Note that this step does not require shuffling per se. This step can also occur by systematically choosing every original shuffle/columns of MaxSums to compare against the rest of MaxSums rather than randomly reselecting the shuffles/columns. This alternate method was tested and did not considerably affect the performance of the method as compared to the results using an additional shuffling that we present here.

The same procedure is then carried out on the real, unshuffled data one time. The best value of adjacent test statistic sums for each window-size are noted along with the location of the best window. As with the randomly chosen shuffled data, these values are compared against MaxSums, produced during the shuffling, to generate raw p-values for each window size. These raw p-values are then compared against the distribution of best raw p-values determined during the second shuffling procedure, to provide a final, protected p-value. This is given by the proportion of best raw p-values in the distribution that are less than or equal to the given raw p-value. This final p-value can be determined as the probability that the best window across all sizes in a randomized dataset will have an unprotected p-value as low as what was observed for the corresponding window size at its best location in the unshuffled data.

### Simulation procedure

Simulated data consisted of two experimental conditions generated from independent Gaussian noise. A ground-truth effect was introduced by adding a boxcar signal of fixed amplitude to one condition over a contiguous region of the data while leaving the second condition unchanged. The effect amplitude was set to 0.1 relative to the standard deviation of the Gaussian noise, and the effect width was fixed at 15 contiguous samples. The objective was to determine whether each statistical method could correctly identify the presence and location of the underlying effect while appropriately controlling for multiple comparisons. AWSS was compared against conventional cluster-based permutation testing using three different cluster-forming thresholds. Performance was quantified as the proportion of simulated datasets in which each method detected a significant effect at each location along the sampled continuum.

### Application to Human Intracranial Recordings

To demonstrate the application of AWSS to biological data, we applied the method to intracranial recordings obtained from three patients with medically refractory epilepsy undergoing stereo-electroencephalography (sEEG) monitoring. Participants performed a sentence-reading task in which sentences of varying syntactic structures were presented using the Rapid Serial Visual Presentation (RSVP) paradigm, with individual words presented sequentially at a rate of 600 ms per word. Comprehension questions followed a subset of trials to encourage attentive reading.

Neural activity was quantified using high gamma power (HGP; 70–170 Hz), which was averaged across the high gamma frequency band and expressed in units of decibels relative to baseline. For the present demonstration, HGP time courses were compared between function words and semantic content words recorded from electrodes located within the anterior temporal lobe. AWSS was applied to identify contiguous temporal windows exhibiting statistically significant differences between the two conditions while correcting for multiple comparisons across all possible window sizes.

## Results

We generated simulated data with a known ground truth effect by generating random noise and adding a box car bolus of a specified amplitude and width to the randomly generated noise. We then tested AWSS on this data along with other existing methods in the field. Cluster refers to the technique described in Maris & Oostenveld (2007) that we tested here with 3 different threshold parameter levels. We intend to expand our results to include TFCE as a comparison as well. For this parameter combination with a low, broad underlying effect, AWSS outperforms the other methods.

Figure 2 illustrates the performance of AWSS and conventional cluster-based permutation testing for one representative simulation parameter combination. The simulated datasets contained a broad, low-amplitude ground-truth effect with an amplitude of 0.1 standard deviations extending across 15 contiguous samples.

**Figure 2.**
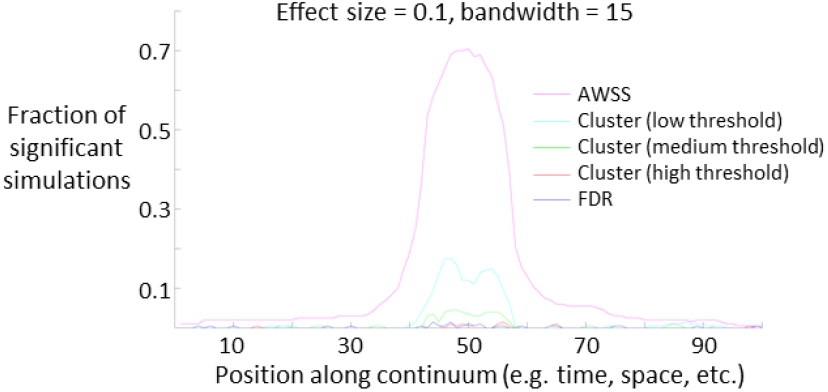
Simulation results. We generated simulated data with a known ground truth effect by generating random noise and adding a box car bolus of a specified amplitude and width to the randomly generated noise. We then tested AWSS on this data along with other existing methods in the field. Cluster refers to the technique described in Maris, E., & Oostenveld, R. (2007) that we tested here with 3 different threshold parameter levels. We intend to expand our results to include TFCE as a comparison as well. For this parameter combination with a low, broad underlying effect, AWSS outperforms the other methods.

AWSS demonstrated substantially greater sensitivity than all cluster-based methods tested for this parameter combination. The principal tradeoff was the appearance of modest “sidelobes,” in which the detected significant region extended somewhat beyond the true boundaries of the simulated effect. This behavior reflects the fact that neighboring windows containing portions of the true signal also accumulate substantial evidence for significance. Future work will systematically evaluate performance across a broader range of effect amplitudes, effect widths, noise structures, and real neurophysiological datasets, as well as compare AWSS with threshold-free cluster enhancement (TFCE).

### Application to Human Intracranial Recordings

To demonstrate the applicability of AWSS to real biological data, we analyzed HGP responses recorded from three representative anterior temporal lobe electrodes, each obtained from a different participant. Across all three recordings (**Figure 3**), AWSS identified significant temporal intervals during which responses to semantic content words exceeded responses to function words. The significant windows corresponded closely to periods of elevated HGP associated with lexical-semantic processing while providing precise temporal localization of the detected effects. These analyses illustrate the practical utility of AWSS for identifying statistically significant neural responses whose temporal extent is not known a priori.

**Figure 3.**
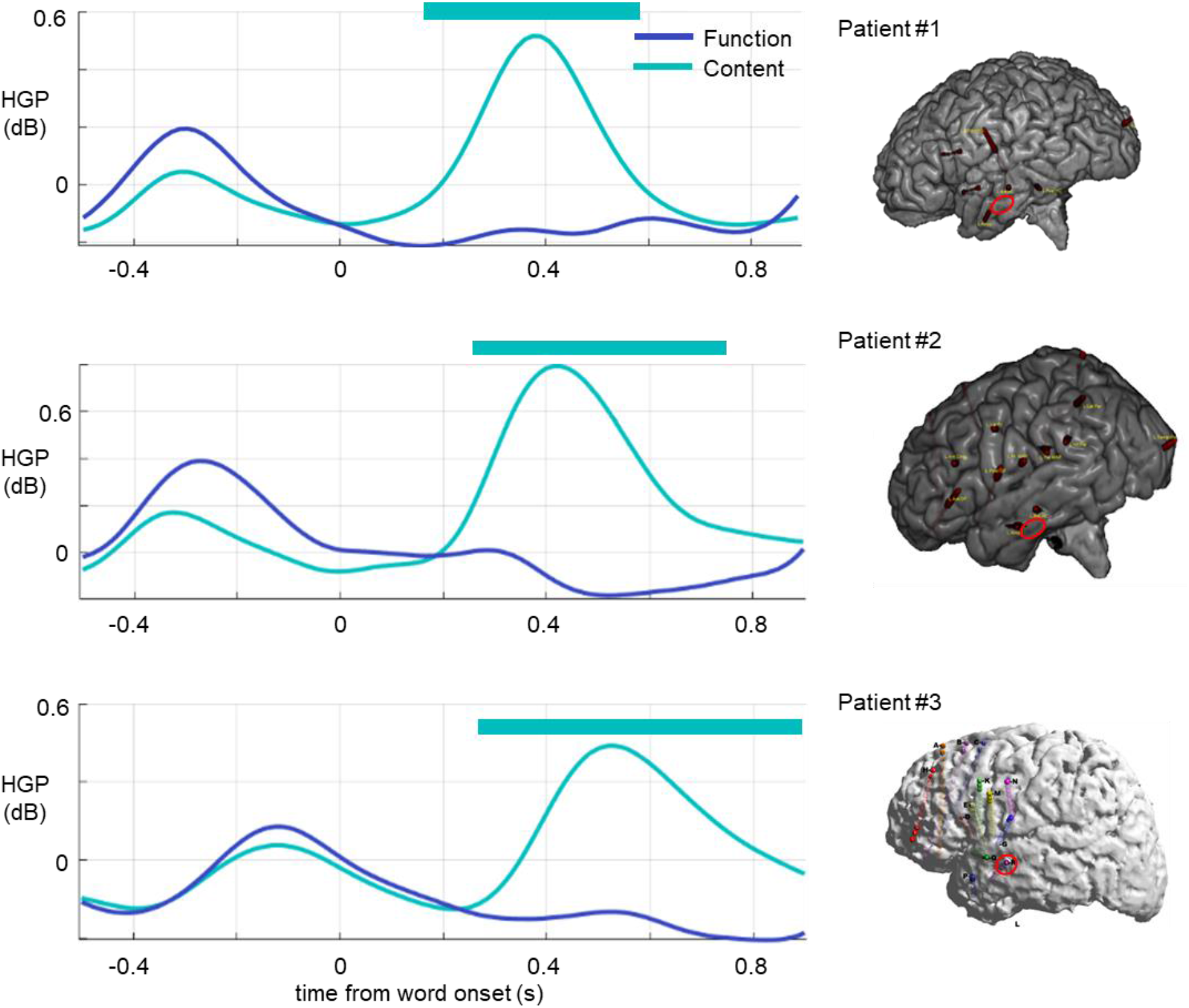
Application of the All Window-Size Search method to human intracranial recordings. High gamma power (HGP; 70–170 Hz) averaged across the high gamma frequency band and expressed in units of decibels is shown for representative electrodes from three different participants (top, middle, and bottom panels). All electrodes were located within the anterior temporal lobe. Dark blue and light blue traces correspond to function words and semantic content words, respectively. Shaded regions (or significance markers, depending on your figure) indicate temporal windows identified by AWSS as exhibiting statistically significant differences between the two conditions after correcting for multiple comparisons across all possible window sizes.

## Discussion

The present work introduces the All Window-Size Search (AWSS) method, a permutation-based framework that controls the family-wise error rate while adaptively searching across all contiguous window sizes. Rather than requiring the investigator to specify the scale of the effect in advance, AWSS evaluates every possible window size and statistically accounts for this adaptive search process. In the simulation presented here, this approach yielded improved sensitivity for detecting broad, low-amplitude effects relative to conventional cluster-based permutation methods. We further demonstrated the applicability of AWSS to real human intracranial neurophysiological recordings, illustrating its practical utility for analyzing biological data.

A central conceptual contribution of AWSS is that it formalizes the statistical search process that researchers naturally perform when visually inspecting ordered data. When examining time series, spatial measurements, or other ordered data, experienced investigators instinctively evaluate effects over multiple temporal or spatial scales. Small, focal effects and broad, diffuse effects are both considered simultaneously, even if this process is not explicitly acknowledged. Conventional statistical analyses generally require investigators to predefine the scale at which testing will occur or rely on methods whose performance depends on assumptions about cluster formation. AWSS instead provides a formal statistical framework that directly accounts for this multiscale search while maintaining family-wise error control.

Unlike conventional cluster-based permutation methods, AWSS is not dependent on an arbitrary cluster-forming threshold. Cluster methods require selecting an initial threshold that determines which neighboring samples become part of a cluster, and the sensitivity of the analysis can vary depending on this choice. Although commonly used thresholds are often reasonable, they remain user-defined analysis parameters. By avoiding threshold-dependent cluster formation, AWSS eliminates one source of investigator degrees of freedom and provides a single, unified framework that simultaneously evaluates all contiguous window sizes.

An additional advantage of AWSS is its generality. The framework is independent of the specific test statistic used to evaluate each sample and therefore can be combined with a wide variety of statistical tests, provided that an appropriate permutation procedure can be defined.

Consequently, the method has potential applications well beyond the motivating examples considered here, including electrophysiology, neuroimaging, behavioral time series, genomics, and other domains involving ordered measurements.

In addition to simulation studies, we demonstrated the applicability of AWSS to real human intracranial neurophysiological recordings. These analyses illustrate that the method can successfully identify statistically significant effects in biological datasets whose temporal extent is not known a priori.

The present simulations examine one representative parameter combination. Future work will systematically characterize performance across a broader range of effect amplitudes, widths, noise structures, and correlation structures.

The present study represents an initial proof of concept. The current implementation focuses on one-dimensional ordered data, but the underlying principles naturally extend to multidimensional spatial and spatiotemporal datasets. Extending the framework to two- and three-dimensional searches may provide a useful approach for applications such as volumetric neuroimaging, cortical surface analyses, and other high-dimensional biological datasets. Future work will also include more comprehensive simulation studies across a broader range of effect sizes, effect widths, noise structures, and comparisons with additional methods, including threshold-free cluster enhancement (TFCE). More broadly, AWSS provides a statistical framework for detecting effects of unknown extent by explicitly correcting the adaptive multiscale search that investigators inevitably perform, thereby increasing sensitivity while maintaining rigorous control of the family-wise error rate.

## References

Benjamini, Y., & Hochberg, Y. (1995). Controlling the False Discovery Rate: A Practical and Powerful Approach to Multiple Testing. Journal of the Royal Statistical Society: Series B (Methodological), 57(1), 289–300. 10.1111/j.2517-6161.1995.tb02031.x

Maris, E., & Oostenveld, R. (2007). Nonparametric statistical testing of EEG- and MEG-data. Journal of Neuroscience Methods, 164(1), 177–190. 10.1016/j.jneumeth.2007.03.024

Smith, S. M., & Nichols, T. E. (2009). Threshold-free cluster enhancement: Addressing problems of smoothing, threshold dependence and localisation in cluster inference. NeuroImage, 44(1), 83–98. 10.1016/j.neuroimage.2008.03.061

